# Megadalton-sized dityrosine aggregates of α-synuclein retain high degrees of structural disorder and internal dynamics

**DOI:** 10.1101/2020.07.26.202358

**Authors:** Silvia Verzini, Maliah Shah, Francois-Xavier Theillet, Adam Belsom, Jan Bieschke, Erich E. Wanker, Juri Rappsilber, Andres Binolfi, Philipp Selenko

## Abstract

Heterogeneous aggregates of the human protein α-synuclein (αSyn) are abundantly found in Lewy body inclusions of Parkinson’s disease patients. While structural information on classical αSyn amyloid fibrils is available, little is known about the conformational properties of disease-relevant, non-canonical aggregates. Here, we analyze the structural and dynamic properties of megadalton-sized dityrosine adducts of αSyn that form in the presence of reactive oxygen species and cytochrome *c*, a proapoptotic peroxidase that is released from mitochondria during sustained oxidative stress. In contrast to canonical cross-β amyloids, these aggregates retain high degrees of internal dynamics, which enables their characterization by solution-state NMR spectroscopy. We find that intermolecular dityrosine crosslinks restrict αSyn motions only locally whereas large segments of concatenated molecules remain flexible and disordered. Indistinguishable aggregates form in crowded *in vitro* solutions and in complex environments of mammalian cell lysates, where relative amounts of free reactive oxygen species rather than cytochrome *c* are rate limiting. We further establish that dityrosine adducts inhibit classical amyloid formation by maintaining αSyn in its monomeric form and that they are non-cytotoxic despite retaining basic membrane-binding properties. Our results suggest that oxidative αSyn aggregation scavenges cytochrome *c*’s activity into the formation of amorphous, high molecular-weight structures that may contribute to aggregate diversity in Lewy body deposits.

## Introduction

Lewy body (LB) aggregates of the pre-synaptic protein αSyn are pathological hallmarks of Parkinson’s disease (1, 2) and related synucleinopathies (3). Whereas ageing and cellular oxidative stress are common denominators in synucleinopathies, LB morphologies and αSyn aggregate/fibril structures differ in a source-of-origin and disease-specific manner (4-7), suggesting that aggregation pathways, fibril strains and aggregate compositions are likely influenced by context-dependent factors that are altogether poorly understood (8). Similarly, the toxicity and spreading of αSyn oligomers and fibrils varies between different brain regions and cell types (9), which underscores the confounding compositional, functional and structural heterogeneity of αSyn aggregates in these disorders (10).

Mitochondria emerged as key organelles in mediating possible aggregation scenarios of αSyn (11), especially in relation to reactive oxygen species (ROS) production, oxidative stress sensing, and as executioners of programmed cell death, i.e. apoptosis (12). Recent findings substantiate this notion by revealing direct interactions between αSyn and mitochondrial membranes and lipids (13, 14), αSyn-induced organelle toxicity (15, 16,Mahul-Mellier, 2020 #93), stress-related re-localization of αSyn to mitochondria (17, 18) and pathological co-localization with mitochondria (and other non-proteinaceous structures) in brain-derived LB inclusions (5) and cellular models of LB formation and maturation (4). In addition, the functions of many other Parkinson’s disease proteins converge on mitochondria and are directly linked to mitochondrial biology (19).

Cytochrome *c* (cyt *c*) resides in the inner mitochondrial membrane (IMM) where it is bound to cardiolipin, an organelle-specific, negatively-charged phospholipid (20). Under physiological cell conditions, cyt *c* transfers electrons between complex III and IV of the electron transport chain and oxidizes superoxide anions to molecular oxygen (O_2_) as a ROS scavenger. Upon cumulating oxidative stress, cardiolipin oxidation frees cyt *c* from the IMM and the protein is actively released into the cytoplasm where it acts as an apoptotic messenger (21). Cytoplasmic cyt *c* ultimately induces caspase activation and cell death, although mitochondrial outer membrane permeation (MOMP) and cyt *c* release do not strictly trigger apoptosis and several of non-deleterious functions have been discovered recently (21). Amongst them, cyt *c* acts as a peroxidase and oxidizes proteins and lipids in the presence of ROS (22). In substrates such as αSyn, cyt *c*-mediated oxidation primarily targets exposed tyrosine residues and leads to the formation of dityrosine crosslinks that locally concatenate individual molecules into covalent, high molecular weight aggregates (23, 24).

Dityrosine adducts of αSyn are found in Lewy body deposits of Parkinson’s disease (PD) patients and in neuronal inclusions of PD mouse models (24, 25) that stain positive for cyt *c* (26, 27). Cyt *c* mediated dityrosine aggregation of αSyn has been reconstituted with isolated components *in vitro* (27-33), however, no attempts have been made to study the structures of the resulting species at the atomic-resolution level. Here, we delineate the structural and dynamic properties of αSyn dityrosine aggregates by solution-state nuclear magnetic resonance (NMR) spectroscopy and complementary methods including dynamic light scattering (DLS), circular dichroism (CD), atomic force (AFM) and negative-stain transmission electron microscopy (TEM), and mass spectrometry (MS). We show that despite their exceedingly large size, they retain individually crosslinked αSyn molecules in highly dynamic, disordered conformations that resemble the biophysical properties of the monomeric protein. Importantly, we establish that residues of the amyloidogenic NAC region of αSyn are minimally perturbed in these assemblies and display no signs of β-aggregation, regardless of their close proximity and high local concentrations. We further demonstrate that cyt *c*-mediated αSyn aggregates form in macromolecularly crowded, cellular environments, where they exhibit structural and dynamic characteristics that are indistinguishable from aggregates reconstituted with purified components *in vitro*.

## Results

### αSyn dityrosine crosslinking produces high molecular weight aggregates

To obtain structural insights into αSyn dityrosine adducts, we reconstituted peroxidase reactions with equal amounts of recombinant, N- terminally acetylated αSyn (50 μM) and purified cytochrome *c* (cyt *c*) to which we added 50, 100, or 500 μM of hydrogen peroxide (H_2_O_2_). We resolved the resulting mixtures by denaturing sodium dodecyl sulfate polyacrylamide gel electrophoresis (SDS-PAGE) and visualized proteins by Coomassie staining (**Figure 1A-C**). In line with previous reports (23, 27, 28), we detected multiple low molecular weight aggregate (LMWAs) species at limiting H_2_O_2_ concentrations, whereas high molecular weight aggregates (HMWAs) formed in the presence of excess peroxide (**Figure 1A**). Although we did not observe protein precipitation in any of these reaction mixtures, soluble HMWAs were retained in the loading slots and stack portions of respective SDS gels. At high H_2_O_2_ concentrations, αSyn was quantitatively incorporated into HMWAs, whereas input levels of monomeric cyt *c* were largely preserved, as expected for an enzymatic reaction and in agreement with published data (28). To evaluate the minimal stoichiometric requirements for cyt *c* in these reactions, we reconstituted oxidative αSyn aggregation with reduced amounts of peroxidase (**Figure 1B**). At αSyn (50 μM) to cyt *c* molar ratios of 100:1 and 50:1, LMWAs formed readily. At a 10:1 ratio of αSyn to cyt *c*, most of αSyn was incorporated into HMWAs. These results confirmed that aggregation of αSyn into covalent adducts in the presence of H_2_O_2_ only required catalytic amounts cyt *c*. Next, we asked whether LMWAs constituted reaction intermediates *en route* to HMWA formation. We added increasing amounts of cyt *c* to preformed LMWAs and allowed reactions to pursue for 30 min (**Figure 1C**). Having resolved the respective mixtures by SDS-PAGE, we found that LMWAs efficiently converted into HMWAs and that final HMWA abundance correlated with cyt *c* input concentrations.

**Figure 1:**
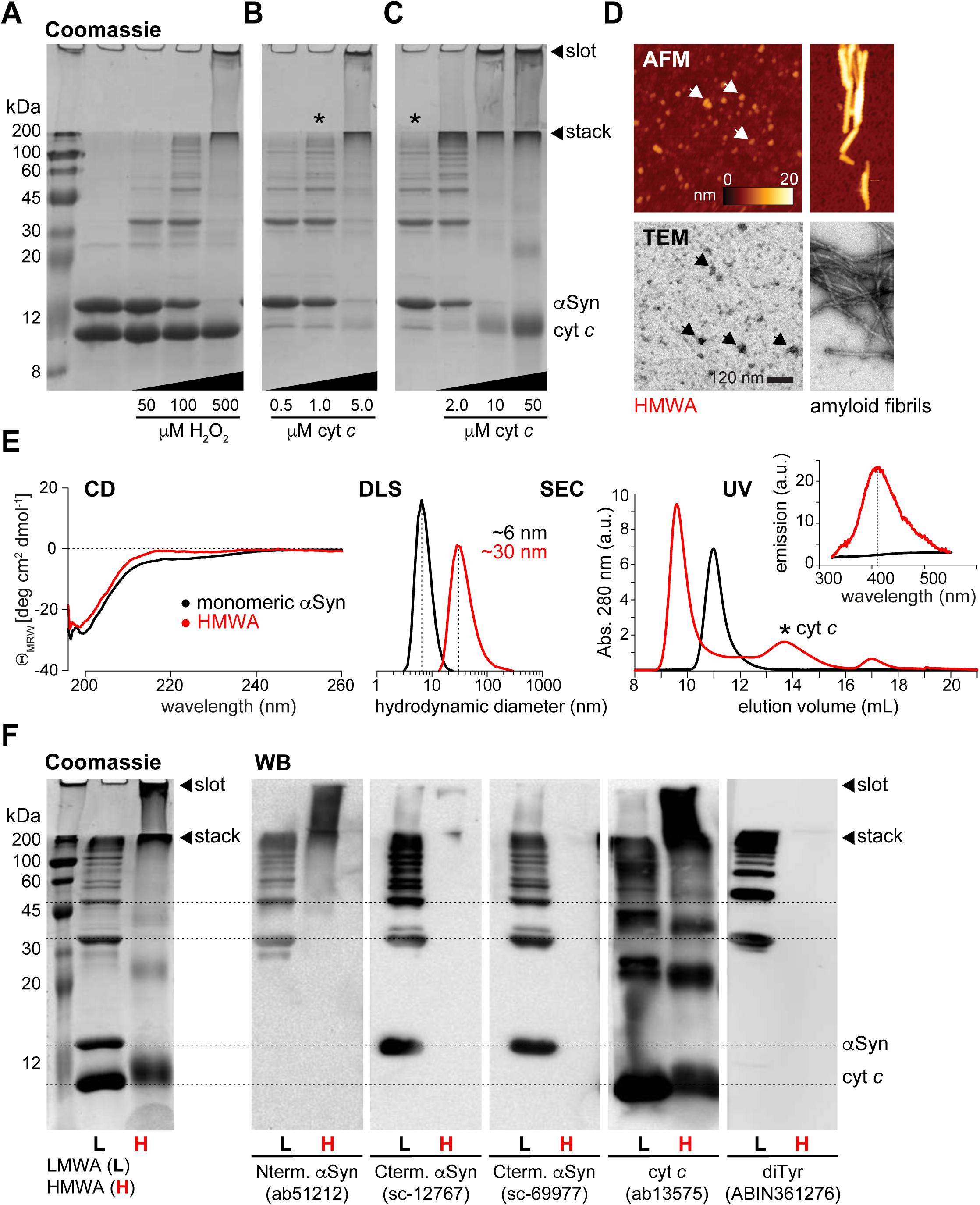
Biochemical and biophysical characterization of dityrosine αSyn aggregates. **(A)** Coomassie-stained SDS-PAGE of low and high molecular weight aggregates of αSyn (50 μM) with 50 μM cyt *c* and 50, 100 or 500 μM H_2_O_2_. **(B)** Aggregation of αSyn (50 μM) with 0.1, 1 or 5 μM cyt *c* and 10 mM H_2_O_2_. **(C)** Preformed low molecular weight aggregates (50 μM αSyn, 1 μM cyt *c*, 10 mM H_2_O_2_, marked with an asterisk) reacted with 2, 10 or 50 μM cyt *c*. All reactions were allowed to proceed for 30 min at 25 °C. Loading slots and gel stacks are indicated with arrowheads. **(D)** Atomic force microscopy (AFM, top) and negative stain transmission electron microscopy (TEM, bottom) of purified high molecular weight aggregates (HMWAs) of αSyn. Control amyloid fibrils are shown on the right. **(E)** Biophysical characterization of monomeric αSyn (50 μM, black) and purified HMWAs (50 μM, red) by CD spectroscopy, DLS, SEC and fluorescence emission spectroscopy. **(F)** Coomassie-stained SDS-PAGE of αSyn low and high molecular weight aggregates (LMWAs and HMWAs, left) and corresponding Western blot (WB) analyses, with antibodies against N- and C-terminal αSyn epitopes, cyt *c* and dityrosine adducts. Dotted horizontal lines connect WB signals to corresponding Coomassie protein bands. Loading slots and gel stacks are indicated with arrowheads.

Aiming to further characterize the structural details of cyt c/H_2_O_2_-mediated αSyn aggregates, we purified HMWAs and analyzed them by AFM and negative-stain TEM (**Figure 1D**). Both methods revealed amorphous globular structures of 30-50 nm diameters (10-15 nm heights), with different macroscopic features compared to canonical amyloid fibrils. Solution characterization by CD spectroscopy, DLS, size-exclusion chromatography (SEC) and ultraviolet (UV) absorption measurements showed that HMWAs retained high degrees of structural disorder and uniform DLS distributions centered at ∼30 nm, roughly the diameter of assembled eukaryotic ribosomes (34), as compared to ∼6 nm for monomeric αSyn (**Figure 1E**). HNWAs eluted as single peaks in SEC void volumes, separated from non-incorporated cyt *c* and well set off from monomeric αSyn, and displayed UV emission bands at 404 nm, characteristic for dityrosine adducts (35). To determine the molecular compositions of αSyn aggregates, we probed LMWAs and HMWAs with different αSyn, cyt *c* and dityrosine-specific antibodies by Western blotting (**Figure 1F**). Antibodies against N-terminal (aa6-23), or C-terminal (aa120-125) αSyn epitopes recognized dimer-, trimer- and higher oligomeric-species in LMWAs that corresponded in size to Coomassie-stained protein bands, confirming that these adducts contained αSyn. The dityrosine antibody reacted with the same set of bands, validating the presence of intermolecular crosslinks. It showed no signal for monomeric αSyn, suggesting that intramolecular dityrosines were not formed. By contrast, the cyt *c* antibody recognized different sets of low molecular weight bands that corresponded to monomeric cyt *c* and to cyt *c* oligomers with no corresponding Coomassie bands. These adducts contained no αSyn based on Western blotting results, which indicated that αSyn-αSyn dityrosine adducts constituted the primary species in LMWAs. Antibodies against C-terminal portions of αSyn and dityrosines failed to recognize epitopes in HMWAs, arguing for the inaccessibility of respective binding sites. Antibodies against the N-terminus of αSyn and against cyt *c* produced HMWA signals, indicating that N-terminal αSyn residues remained accessible and that some co-aggregated cyt *c* was present. These results confirmed that intermolecular dityrosine crosslinks between individual αSyn molecules formed the basis of low and high molecular weight aggregates.

### Dityrosine aggregates are disordered and dynamic

To derive high-resolution insights into the structural and dynamic properties of αSyn in HMWAs, we reconstituted aggregates with ^15^N isotope-labeled, N-terminally acetylated αSyn and unlabeled cyt *c* in the presence of peroxide. We removed H_2_O_2_ and free cyt *c* by SEC and pooled HMWA fractions for solution-state NMR experiments. Despite the exceedingly large size of these assemblies, usually beyond the scope of solution NMR measurements, 2D ^1^H-^15^N heteronuclear correlation spectra of HMWAs were of excellent quality and revealed the majority of αSyn signals at resonance positions corresponding to those of the monomeric protein (**Figure 2A**). These NMR characteristics substantiated CD results about the disordered nature of αSyn in these aggregates (**Figure 1E**) and established that crosslinked αSyn molecules retained high degrees of internal dynamics. Upon closer inspection of NMR spectra, we noted pronounced chemical shift changes for N-terminal αSyn residues 1-10, including M1 and M5, that we and others had previously identified as characteristic for oxidized methionines (i.e. methionine sulfoxides) (36, 37), confirming that NMR samples had indeed formed under oxidative conditions. Importantly, we also noted continuous stretches of severely line-broadened NMR signals for residues around Y39, Y125, Y133 and Y136, the four tyrosines of αSyn and possible sites of dityrosine crosslinks. To verify that detected NMR signals indeed originated from αSyn aggregates, we performed diffusion ordered spectroscopy (DOSY) measurements on HMWAs (**Figure 2B**). DOSY results indicated that HMWA diffusion was greatly reduced compared to monomeric αSyn and corresponded to a megadalton-sized ∼35 nm diameter particle, in good agreement with TEM and DLS results (**Figure 1D** and **E**). To further explore αSyn dynamics in HMWAs, we analyzed signal intensity attenuations on a residue-resolved basis and performed longitudinal (*R*_*1*_) and transverse (*R*_*2*_) relaxation experiments as well as hetero-nuclear Overhauser effect (hetNOE) measurements (**Figure 2C**). We determined intensity ratios (I/I_0_) for well-resolved HMWA signals (I) in relation to monomeric αSyn (I_0_) and found that site-selective line broadening centered at Y39 and C-terminal Y125, Y133 and Y136. In relation to these sites, attenuations weakened in a distance-dependent manner, with more expansive effects around Y39 compared to Y125, Y133 and Y136 (**Figure 2C**, top). αSyn residues comprising the aggregation-prone non-amyloid component (NAC) region (aa60-95) displayed minimal line-broadening, which suggested that their overall structural and dynamic properties were retained in HMWAs. N-terminal αSyn residues 1-10 exhibited similarly small line-broadening effects, indicating equally preserved dynamics in HMWAs and probably explaining the observed antibody accessibility (**Figure 1F**). Quantitative *R*_*1*_ and *R*_*2*_ results supported these conclusions by establishing that fast ps to ns amide backbone motions (*R*_*1*_) were marginally affected, whereas *R*_*2*_ profiles showed position-dependent attenuations consistent with restricted motions and/or conformational exchange on the μs to ms timescale (**Figure 2C**, middle). hetNOE data confirmed that HMWAs displayed independent segmental motions characteristic for a disordered protein (**Figure 2C**, bottom). The qualitative picture that emerged from these experiments stipulated that dityrosine crosslinks restricted backbone motions of concatenated αSyn molecules only in the vicinity of crosslinked residues, whereas remote sites retained high degrees of flexibility and conformational dynamics. On the macroscopic level, these results supported a model of loosely packed, macromolecular assemblies of disordered molecules held together by repeating, intermolecular side-chain crosslinks.

**Figure 2:**
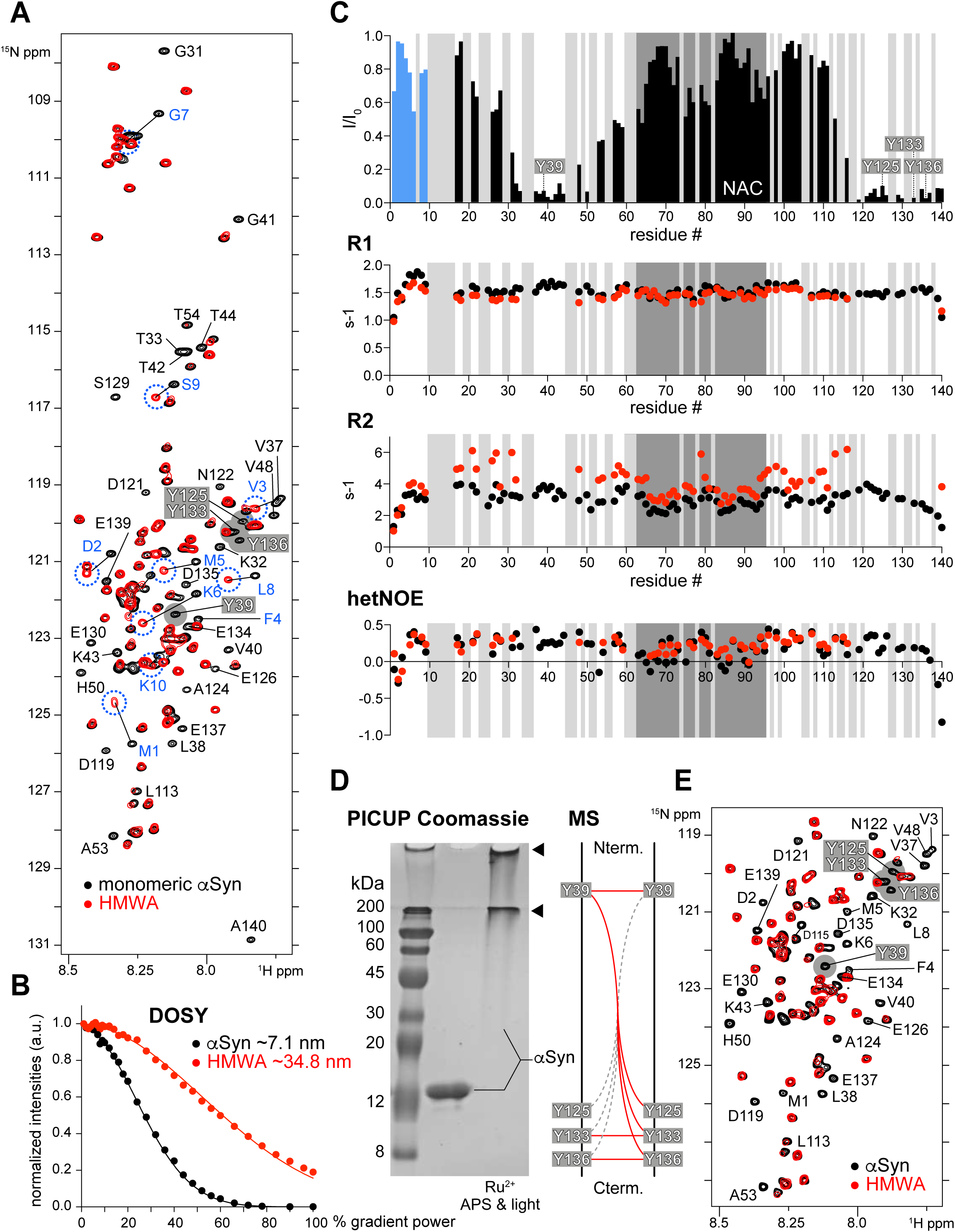
Structural and dynamic characterization of HMWAs. **(A)** Overlay of 2D ^1^H-^15^N NMR spectra of ^15^N isotope-labeled, N-terminally acetylated, monomeric αSyn (50 μM, black) and of HMWAs (50 μM, red). HMWA resonances broadened beyond detection are labeled. Chemical shift changes of N-terminal αSyn residues 1-10 are shown in blue. NMR signals Y39, Y125, Y133 and Y136 are highlighted in grey. **(B)** Diffusion ordered spectroscopy (DOSY) measurements of monomeric αSyn (black) and HMWAs (red). **(C)** *top*: Residue-resolved analysis of NMR signal intensity ratios (I/I_0_) between HMWAs (I) and monomeric αSyn (I_0_). Blue bars correspond to N-terminal αSyn residues 1-10. Black bars denote I/I_0_ values of well-resolved HMWA resonances. *middle:* Overlay of backbone amide R1 (1/T_1_, top) and R2 (1/T_2_, bottom) relaxation rates of monomeric αSyn (black) and HMWAs (red). *bottom:* heteronuclear NOE values of monomeric αSyn (black) and HMWAs (red). Light grey boxes indicate resonances not included in the analysis due to signal overlap. The central NAC region is shown in dark grey. N- and C-terminal tyrosines are highlighted. **(D)** Coomassie-stained SDS-PAGE of αSyn HMWAs (50 μM) generated by photo-induced cross-linking of unmodified proteins (PICUP). Loading slots and gel stacks are indicated by arrows. Summary of dityrosine crosslinks identified in PICUP HMWAs by mass spectrometry (MS). **(E)** Selected region of 2D ^1^H-^15^N NMR spectra of ^15^N isotope-labeled, N-terminally acetylated, monomeric αSyn (50 μM, black) and of PICUP-HMWAs (50 μM, red). HMWA resonances broadened beyond detection are labeled, NMR signals of Y39, Y125, Y133 and Y136 are highlighted in grey. All NMR spectra were recorded at 283 K (10 °C).

To independently verify the general nature of such dityrosine aggregate characteristics, we resorted to photoinduced cross-linking of unmodified proteins (PICUP) and reacted N-terminally acetylated αSyn with ammonium-persulfate (APS) and Ruthenium (Ru^3+^) under light as reported previously (29). PICUP HMWAs displayed similar SDS-PAGE migration properties as cyt *c*/H_2_O_2_-mediated aggregates in that αSyn was quantitatively converted into high molecular weight species retained in loading slots and stacking gels (**Figure 2D**). Exploiting the homogenous nature of PICUP HMWAs, we analyzed aggregates by MS to delineate dityrosine positions and relative distributions (**Figure S1A**). We identified crosslinks of Y39 peptides to αSyn fragments containing Y39, Y125, Y133 and Y136. Similarly, we detected intermolecular Y133-Y133 and Y136-136 crosslinks, but no intra- or inter-molecular adducts between other C-terminal tyrosines (**Figure 2D** and **Figure S1A**). These results supported the central role of Y39 as a connectivity hub in αSyn PICUP aggregates, in line with previous findings (29). To compare the structural features of PICUP HMWAs and cyt *c*/H_2_O_2_-mediated assemblies, we recorded NMR spectra on photo-induced aggregates of ^15^N isotope-labeled αSyn (**Figure 2E** and **Figure S1B**). Surprisingly, both types of HMWAs displayed virtually indistinguishable spectral features, especially in terms of site-selective line broadening. Different to cyt *c*/H_2_O_2_ HMWAs, however, we did not observe N-terminal αSyn residues 1-10 in NMR spectra of PICUP aggregates. Our combined NMR data established that dityrosine HMWAs of αSyn exhibited similar structural and dynamic properties, irrespective of whether they were generated via a protein-catalyzed peroxidase reaction or directly by photo-induced crosslinking.

### Dityrosine aggregates form in complex environments

Next, we asked whether cyt *c*/H_2_O_2_ HMWAs of αSyn formed in the presence of competing interactions with other macromolecules in crowded environments. We initially employed bovine serum albumin (BSA), which has been shown to transiently interact with N-terminal αSyn residues and to bind to Y39 via weak hydrophobic contacts (18, 38), and reconstituted cyt *c*/H_2_O_2_ aggregation reactions with 15N isotope-labeled αSyn and increasing amounts of unlabeled BSA (**Figure 3A**). SDS-PAGE analysis revealed that stacking gel-retained HMWAs specifically formed in solutions of up to 10 mg/mL of BSA, which was the highest concentration we used to avoid overloading the gel. We reacted ^15^N isotope-labeled αSyn with unlabeled cyt *c* and H_2_O_2_ in the presence of a 20-fold higher amount of BSA (200 mg/mL) and recorded *in situ* NMR experiments on the resulting mixture (**Figure 3B** and **Figure S2A**). Similar to previous results, we obtained high quality NMR spectra that revealed the characteristic line-broadening profile of dityrosine-crosslinked αSyn, which confirmed that cyt *c*-H_2_O_2_ aggregation of αSyn occurred under highly crowded *in vitro* conditions and regardless of protein-protein interactions that involved Y39. Following these observations, we extended our analysis to neuronal RCSN3 cell lysates (39). We adjusted total lysate protein concentrations to 3 mg/mL and added equimolar amounts of αSyn and cyt *c* (50 μM) and 0.5 - 500 mM of H_2_O_2_. Western blotting revealed that aggregate formation only occurred at the highest peroxide concentration (**Figure 3C**, top), in contrast to previous experiments with isolated components (**Figure 1**) and in BSA-crowded solutions (**Figure 3A**). We hypothesized that peroxide-metabolizing enzymes may have cleared exogenously added H_2_O_2_. Indeed, when we measured remaining H_2_O_2_ levels using a colorimetric assay, we found that only 1.8 ± 0.8% of input peroxide (10 mM) was present in these lysates (**Figure 3C**). To weaken endogenous peroxidase activities, we treated RCSN3 lysates with diethyl pyrocarbonate (DEPC), a chemical that inactivates enzymes by irreversible carbethoxylation of active site residues (40). Because DEPC has a limited lifetime in aqueous solutions and decomposes readily, one of its advantages over enzyme inactivation by acid, SDS, or high temperature denaturation is the perseveration of otherwise native lysate conditions and the absence of chaotropic species after treatment. Accordingly, we adjusted the timing of lysate inactivation by DEPC (**Figure S2B**) and added αSyn, cyt *c* and H_2_O_2_ as outlined before (**Figure 3C**, bottom). Under these conditions, 40 ± % of exogenously added peroxide (10 mM) was preserved and we detected αSyn aggregation at correspondingly lower H_2_O_2_ concentrations. We repeated these reactions with ^15^N isotope-labeled αSyn and recorded *in situ* NMR experiments in DEPC-treated RCSN3 lysates (**Figure 3D** and **Figure S2C**). 2D NMR spectra exhibited compelling degrees of similarity with other HMWA samples. We clearly observed the spectral fingerprints of dityrosine crosslinks manifested by the conserved line-broadening of residues neighboring Y39, Y125, Y133 and Y136, and the characteristic chemical shift changes of N-terminal αSyn residues due to methionine oxidation (see **Figure 2A** for comparison). Together with results in BSA-crowded solutions, these findings demonstrated that cyt *c*/H_2_O_2_-mediated aggregates of αSyn formed efficiently in complex physiological environments, especially under conditions of sustained ROS availability required for dityrosine crosslinking.

**Figure 3:**
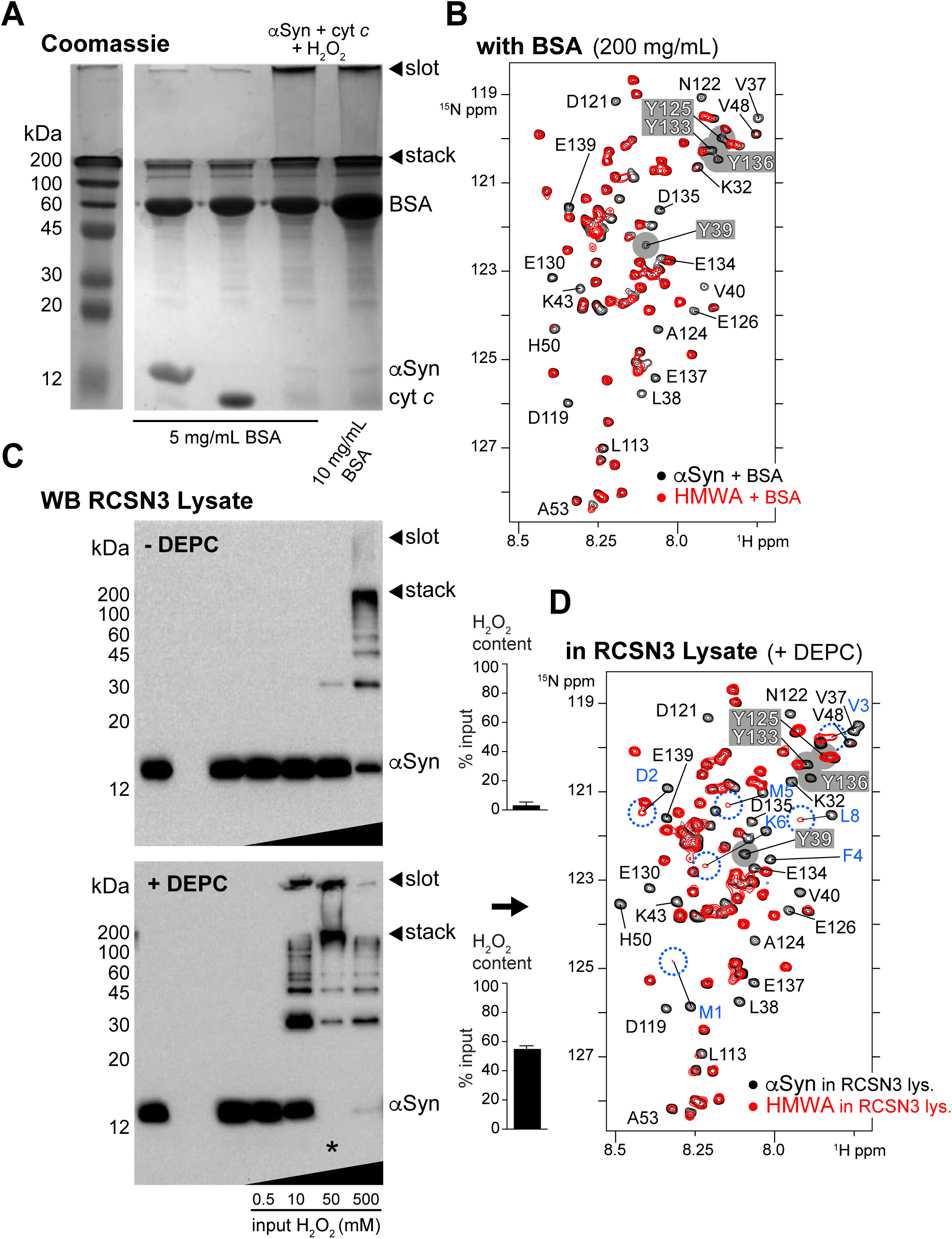
HMWA formation in crowded environments. **(A)** Coomassie-stained SDS-PAGE of αSyn (50 μM) aggregation reactions with cyt *c* (50 μM) and H_2_O_2_ (10 mM) in the presence of 5 or 10 mg/mL of BSA. Loading slots and gel stacks are indicated by arrows. **(B)** Selected region of 2D ^1^H-^15^N NMR spectra of ^15^N isotope-labeled, N-terminally acetylated, monomeric αSyn (50 μM) in the presence of 200 mg/mL BSA (black). Note that N-terminal αSyn residues 1-10 in the reference spectrum are broadened beyond detection due to transient interactions with BSA. Y39 signal intensity of monomeric αSyn is attenuated for the same reason. The corresponding *in situ* NMR spectrum of HMWAs (50 μM) formed in the presence of BSA is overlaid in red. Broadened HMWA resonances are labeled, NMR signals of Y39, Y125, Y133 and Y136 are highlighted in grey. **(C)** HMWA formation in RCSN3 cell lysates (total protein concentration 3 mg/mL). *top:* Western blotting of aggregation reactions in native RCSN3 lysates upon addition of αSyn (50 μM), cyt *c* (50 μM) and 0.5 – 500 mM H_2_O_2_. Colorimetric quantification of remaining peroxide levels after addition of 10 mM H_2_O_2_. *bottom:* Western blotting of aggregation reactions in DEPC-inactivated RCSN3 lysates upon addition of αSyn (50 μM), cyt *c* (50 μM) and 0.5 – 500 mM H_2_O_2_, with remaining peroxide levels after addition of 10 mM H_2_O_2_ shown on the right. **(D)** Selected region of 2D 1H-15N NMR spectra of 15N isotope-labeled, N-terminally acetylated, monomeric αSyn (50 μM) in DEPC-treated RCSN3 lysate (black) and *in situ* NMR spectrum of HMWAs (50 μM) formed after addition of cyt *c* (50 μM) and 50 mM H_2_O_2_ (red). HMWA resonances broadened beyond detection are labeled. Chemical shift changes of N-terminal αSyn residues 1-10 are shown in blue. NMR signals Y39, Y125, Y133 and Y136 are highlighted in grey. All NMR spectra were recorded at 283 K (10 °C).

### Dityrosine aggregates bind membranes, inhibit amyloid formation and are non-cytotoxic

Because N-terminal αSyn residues that mediate lipid-anchoring contacts (41) remained accessible in HMWAs, we asked whether aggregates retained the ability to interact with membranes. We added ^15^N isotope-labeled monomeric and HMWA αSyn to small unilamellar vesicles (SUVs) that we reconstituted from pig-brain polar lipid extract. In line with previous results on such SUVs (38), monomeric αSyn displayed continuous line-broadening of residues 1-100, which serves as a characteristic NMR-indicator for αSyn membrane binding (42, 43) (**Figure 4A** and **Figure S3A**). Surprisingly, 2D NMR spectra of ^15^N isotope-labeled HMWAs exhibited similar degrees of line-broadening in the presence of SUVs (**Figure 4B** and **Figure S3B**). Remaining HMWA resonances included residues within the first 100 amino acids of αSyn, which suggested that HMWA-SUV contacts were different than those of the monomeric protein. Because αSyn interacts with membranes in helical conformations (44), we asked whether HMWAs adopted similar helical structures and recorded CD spectra of monomeric and HMWA αSyn in the presence of SDS micelles, a commonly used surrogate for studying Syn’s α-helical conversion upon membrane binding (45) (**Figure 4C**). We found that monomeric and HMWA αSyn displayed similar helical signatures in corresponding CD spectra, which suggested that HMWA-membrane interactions triggered some degrees of α-helical rearrangements.

**Figure 4:**
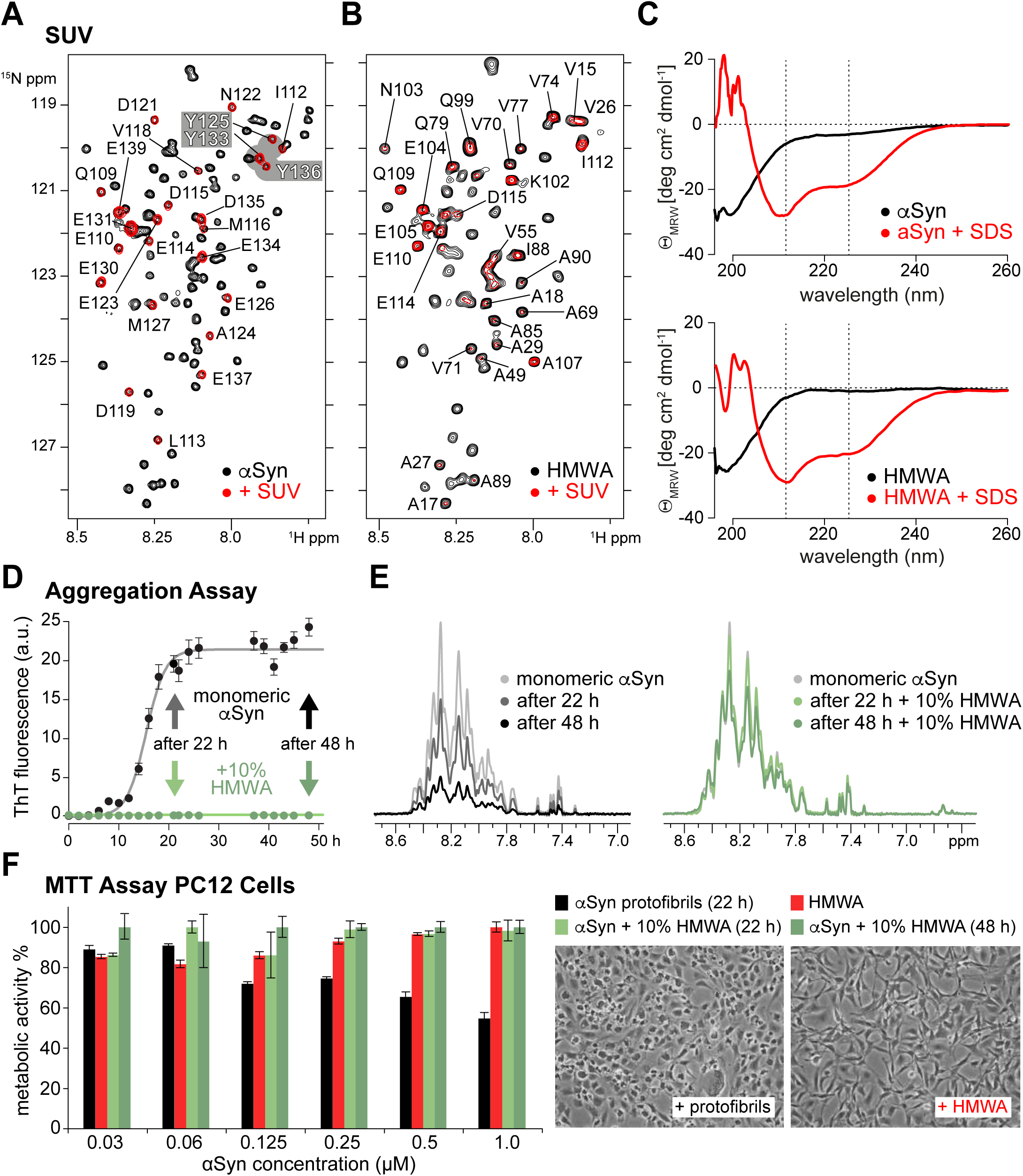
HMWA membrane-binding, aggregation inhibition and cellular toxicity. **(A)** Selected region of 2D ^1^H-^15^N NMR spectra of ^15^N isotope-labeled, N-terminally acetylated, monomeric αSyn (50 μM, black) and in the presence of small unilamellar vesicles (SUV, red). Non-broadened NMR signals of C-terminal αSyn residues are labeled. Y125, Y133 and Y136 are highlighted in grey. **(B)** Same selected region of ^15^N isotope-labeled HMWAs (50 μM, black) and upon binding to SUVs (red). NMR spectra were recorded at 303 K (30 °C). Non-broadened NMR signals are labeled. **(C)** *top:* Overlay of CD spectra of monomeric αSyn (50 μM, black) and in the presence of SDS micelles (red). *bottom:* Overlay of CD spectra of purified HMWAs (50 μM, black) and in the presence of SDS micelles (red). **(D)** Kinetic β-aggregation assays and measured Thioflavin-T (ThT) fluorescence emission for monomeric, ^15^N isotope-labeled αSyn (50 μM, black) and in the presence of 10% unlabeled HMWAs (green). Error bars depict SEM values from triplicate measurements. NMR samples were collected at 22 and 48 h timepoints for respective 1D NMR measurements. **(E)** 1D ^15^N-edited NMR signals of monomeric αSyn in aggregation reactions without (grey to black) and with 10% HMWAs (green). **(F)** Metabolic MTT assays on PC12 cells upon addition of αSyn protofibrils (black), purified HMWAs (red) and aliquots of inhibited aggregation reactions after 22 h (light green) and 48 h (dark green), at the indicated concentrations. Bar graphs denote changes in metabolic activity expressed in % compared to untreated cells. Error bars represent SEM from triplicate measurements. Phase-contrast microscopy images of PC12 cells incubated with 1 μM of αSyn protofibrils or 1 μM of HMWAs are shown on the right.

Finally, we sought to revisit HMWA effects on the aggregation behavior(s) of monomeric αSyn. Previous studies suggested that dityrosine adducts of αSyn impaired the formation of cross-β amyloid structures in a concentration-dependent manner (32). In agreement with these findings, we observed the quantitative inhibition of thioflavin T (ThT)-positive aggregate formation in shaking assays when we added 10% HMWAs to monomeric αSyn (**Figure 4D**). To investigate the mode of inhibition, we repeated these assays with ^15^N isotope-labeled, monomeric αSyn and unlabeled HMWAs, and measured 1D ^15^N-edited NMR spectra at different time-points of the reaction. In the absence of HMWAs, we observed the progressive attenuation of NMR signals as expected for the incorporation of monomeric αSyn into growing amyloid fibrils (**Figure 4E**). In the presence of HMWAs, NMR signal intensities of αSyn did not diminish substantially, suggesting that inhibition was mediated by the preservation of the monomeric protein state and not by the generation of ThT-negative, off-pathway aggregates as has been observed for many other inhibitors (46). To assess whether HMWAs were cytotoxic, we measured the metabolic activity of PC12 cells to which we added protofibrils generated from monomeric αSyn, purified HMWAs and aliquots of inhibited aggregation mixtures containing αSyn and HMWAs that we collected after 22 and 48 hours (**Figure 4F**). Addition of protofibrils decreased the conversion of 3-(4,5-dimethylthiazol-2-yl)-2,5-diphenyltetrazoliumbromide (MTT) into formazan in a concentration-dependent manner, in line with previous reports (47) and characteristic for cytotoxic aggregates (48). By contrast, neither purified HMWAs nor αSyn-HMWAs mixtures impaired cell viability and specimens displayed no signs of cellular stress (**Figure 4F**). These results established that exogenously added dityrosine aggregates of αSyn inflicted no adverse effects on the metabolic activity and viability of PC12 cells.

## Discussion

Recent years saw a wealth of high-resolution structural information about αSyn oligomers (49) and amyloid fibrils (50), ranging from *in vitro* reconstituted assemblies to specimens directly derived from post-mortem brains (51). Correlative imaging studies delivered complementary insights into the compositions and morphologies of Lewy bodies and their complex nature in terms of membrane and organelle inclusions, and the abundance of heterogeneous aggregate structures (4, 5, 10). Parallel to these advancements, functional links between ageing, cellular oxidative stress, neurodegeneration and oxidative αSyn modifications have been explored for some time (52). In this context, our results add primary insights into the structure and dynamics of non-canonical αSyn aggregates characterized by covalent dityrosine crosslinks between N- (Y39) and C-terminal (Y125, Y133, Y136) residues. We demonstrate that these aggregates efficiently form under oxidative conditions in the presence of catalytic amounts of a peroxidase enzyme (cytochrome *c*) (**Figure 1**) and that they constitute amorphous, locally concatenated assemblies that retain high degrees of structural disorder and internal dynamics (**Figure 2A-C**). Although we intentionally chose ‘exhaustive’ reaction conditions to produce homogenous endpoint products (i.e. HMWAs), we reason that the structural and dynamic properties that we elucidated are similarly displayed in mixtures of lower molecular-weight aggregates (i.e. LMWAs), which constitute reaction intermediates (**Figure 1C**). Having established that αSyn dityrosine aggregates also form in artificially crowded *in vitro* solutions and cell lysates (**Figure 3**) further strengthens the notion that crosslinking avidity and specificity are well preserved in complex physiological environments and that these aggregates may co-exist with monomeric, oligomeric or fibrillar forms of αSyn in cells. Furthermore, it seems plausible that mixed dityrosine adducts between different αSyn species can form, especially under conditions of sustained oxidative stress and considering the persistent reactivity of unmodified tyrosine residues. Such mixed crosslinks may modify non-covalent αSyn aggregates or stabilize existing αSyn fibrils (23, 53). In this regard, it is interesting to note that Y39, Y125, Y133 and Y136 in structures of reconstituted αSyn fibrils are not part of the ordered amyloid cores and do not contribute to protofilament packing (50), which suggests that they may remain accessible for oxidative crosslinking. Indeed, results from dityrosine-αSyn EM immunogold co-labeling studies on PD patient-derived Lewy bodies confirm the presence of such species in physiological samples (25). Co-aggregation with non-covalent αSyn fibrils may further explain certain aspects of morphological changes during Lewy body maturation, especially at later stages when fibrillar structures convert into bloated spherical inclusions (4). Intriguingly, this transition coincides with the recruitment of mitochondria into Lewy bodies (4), which are abundantly found in mature deposits (5). This puts αSyn, mitochondrial cytochrome *c* and the sites of ROS production in close proximity and it is straightforward to imagine how these encounters may drive or perpetuate the formation of dityrosine aggregates and impede cellular functions such as proteasomal degradation, chaperone-mediated autophagy (54), or illicit inflammatory activation of neighboring cells upon excretion (55). In this regard, results pertaining to the non-cytoxicity of HMWAs when externally applied to PC12 cells ought to be considered with caution (**Figure 4F**). Because standard MTT assays do not provide information about aggregate internalization, we are unable to assess whether they cross the plasma membrane and reach the cytoplasm to cause intracellular effects. Alternatively, monomeric or oligomeric αSyn may directly engage with mitochondria and actively contribute to ROS production, cytochrome *c* release and, in turn, dityrosine aggregate formation (56). It is important to stress, however, that the generation of αSyn dityrosine adducts does not strictly require a peroxidase enzyme (i.e. cytochrome *c*) and that they can form as reaction side-products of tyrosine nitration, by dopamine-mediated autoxidation, and in the presence of oxidized lipids or divalent metals (52). Thus, cellular scenarios in which such aggregates are generated with or without the involvement of mitochondria are manifold.

Dityrosine crosslinks may further render involved residues inert to other post-translational modifications (PTMs) such as nitration (57) and phosphorylation (58). All αSyn tyrosines undergo reversible phosphorylation (59) and PTM cross-talk between Y125 and S129 phosphorylation, as well as M127 oxidation is well established (37, 60). Crosslinks are likely to impair the accessibility of these sites and, thereby, modulates the enzymatic establishment or removal of individual PTMs. With regard to αSyn S129 phosphorylation, the pathological hallmark of Lewy body inclusions (61), it may help to preserve this modification by restricting access of bulky phosphatase holoenzymes (62). Intriguingly, mitochondrial recruitment and phospho-S129 accumulation coincide with Lewy body maturation (4). Similarly, dityrosine adducts and aberrant PTM states may diminish αSyn interactions with other proteins, including the endocytosis GTPase Rab8 (63) and the synaptic vesicle v-SNARE component synaptobrevin-2/VAMP2 (64), which may exert debilitating effects on the physiological functions of αSyn, probably exacerbated by the residual membrane-binding activity of these adducts (**Figure 4B**). They may further affect C-terminal proteolytic processing events (65) and the cellular clearance of αSyn (52). Given that covalent dityrosine crosslinks represent irreversible protein modifications that cannot be undone by cellular enzymes, they impact αSyn’s biology in a persisting manner.

In summary, our results established that oxidative aggregation of αSyn via dityrosine crosslinking imposes minor effects on N-terminal and central portions of the protein, with residues involved in membrane anchoring and of the amyloidogenic NAC region being largely unperturbed, respectively (**Figure 2**). Resulting degrees of conformational freedom appear to allow for structural rearrangements required for helical membrane-binding (**Figure 4B, C**) and to mediate the inhibition of NAC-dependent cross-β amyloid formation (**Figure 4D, E**), probably via capping of growing fibril ends. Because three of the four tyrosines of αSyn are located in the C-terminus of the protein, functional consequences are expected to be more drastic for activities mapped to this region, including the establishment and removal of PTMs and various protein-protein interactions. With regard to the involvement of cytochrome *c*, oxidative dityrosine crosslinking may scavenge the activity of this proapoptotic messenger and, in turn, delay the onset of programmed cell death, as suggested earlier (27). By doing so, the outlined reaction scenarios may perpetuate detrimental conditions of oxidative stress and, thereby, exacerbate cellular insults that contribute to PD pathology.

## Acknowledgments

We thank Dr. Rudi Lurz (MPI fuer Molekulare Genetik, Berlin) for help with TEM experiments and Dr. Peter Schmieder and Monika Beerbaum for excellent maintenance of NMR infrastructure. A.B. and J.R. acknowledge support by the Wellcome Trust (103139 and 108504) and the Deutsche Forschungsgemeinschaft (DFG, German Research Foundation, 329673113). The Wellcome Centre for Cell Biology is supported by core funding from the Wellcome Trust (203149). M.S., J.B. and E.E.W. were supported by grant no. 3-12056 from the Helmholtz-Israel Cooperation in Personalized Medicine and grants from the German Research Foundation (SFB618, SFB740, Neurocure), the European Union (EuroSpin and SynSys) and the Helmholtz Association (MSBN and HelMA). S.V. A.B. and P.S. were funded by the European Research Council (ERC) Consolidator Grant NeuroInCellNMR (647474) to P.S.

## Materials and Methods

### Proteins

Unlabeled or uniform ^15^N or ^15^N/^13^C isotope-labeled, N-terminally acetylated, human alpha-synuclein (αSyn) was expressed and purified under native conditions as previously described (38). Recombinant proteins were resuspended in 20 mM phosphate buffer, 150 mM NaCl at pH 6.4 (NMR buffer). Protein concentrations were determined by UV/VIS absorption measurements at 280 nm (λ) with an extinction coefficient (ε) of 5600 M^-1^ cm^-1^. Purified horse heart cytochrome *c* (cyt c) was obtained from Sigma, bovine serum albumin was purchased from Roth. Protein stock solutions were prepared in NMR buffer.

### Low and high molecular weight aggregates

Low molecular weight aggregates (LMWAs) of αSyn were obtained by mixing 50 μM αSyn with 50 μM cyt *c* and 100 μM hydrogen peroxide (H_2_O_2_) i.e. at a molar ratio of 1:1:2, unless specified otherwise. High molecular weight aggregates (HMWAs) were obtained by mixing 50 μM αSyn with 50 μM cyt *c* and 2 mM H_2_O_2_ (1:1:200), unless specified otherwise. LMWA and HMWA reactions were incubated for 30 min at 25 °C to yield the respective samples. Molar aggregate concentrations were considered equal to αSyn input concentrations in a first approximation. For AFM and TEM experiments and biophysical characterizations (**Figure 1D, E**), NMR measurements (**Figure 2A** and **E**) and testing of membrane binding and aggregation interference (**Figure 4**), HMWAs were purified by size exclusion chromatography on a Superdex-75 column to remove non-incorporated cyt *c* and excess hydrogen peroxide (H_2_O_2_). Pooled HMWA fractions were either used immediately or flash frozen in liquid nitrogen and stored at −80 °C. For NMR experiments, estimates of purified ^15^N isotope-labeled HMWA concentrations were derived based on 1D ^15^N-edited proton NMR spectra in comparison to reference samples of monomeric αSyn at known concentrations. Accordingly, samples were adjusted to 50 μM.

### Polyacrylamide gel electrophoresis (PAGE)

Denaturing sodium dodecylsulfate (SDS) PAGE gels used in this study were prepared on Miniprotean III systems (BioRad) with a constant separating-gel pore size of 18% acrylamide (w/v). Proteins were visualized by gel staining with Coomassie Brilliant Blue (Sigma Aldrich).

### Atomic force microscopy (AFM)

Sheet mica (Nanoworld) was glued to microscope glass slides and 20 μL of 25 μM HMWAs or αSyn amyloid fibrils were adsorbed for 10 min onto freshly cleaved mica (1 x 1 cm), washed with filtered, deionized water (3 × 30 µL) and dried overnight. Reference amyloid fibrils were obtained by shaking 100 μL of 50 μM αSyn in NMR buffer at 200 rpm, 37 °C for 7 days. Fibrils were stored at 4 °C and shortly vortexed before experiments. Amyloid fibril concentrations were considered equal to input amounts of αSyn. Dry AFM images were recorded on a Nanowizard II/Zeiss Axiovert setup (JPK) using intermittent contact mode and FEBS cantilevers (Veeco).

### Negative-stain transmission electron microscopy (TEM)

20 μL of 25 μM HMWAs and αSyn amyloid fibrils were adsorbed onto glow-discharged carbon-coated copper grids for 1 min. Excess liquid was removed with filter paper and grids were washed twice with H_2_O before staining with 2% (w/v) uranyl acetate for 15 s. TEM images were acquired on a Philips CM100 microscope.

### Circular dichroism (CD) spectroscopy

CD measurements were performed on a Jasco J-810 spectropolarimeter at 25 °C with samples at 10 μM. Far-UV CD spectra were collected in NMR buffer using a 0.1 cm path-length cuvette. Five scans were averaged and blank (buffer) scans were subtracted. CD experiments with SDS-micelle (**Figure 4C**) were performed in the same manner. Micelles were prepared by dissolving 10 mM SDS in NMR buffer as reported in (45).

### Dynamic light scattering (DLS)

DLS measurements were acquired on a Zetasizer Nano ZS (Malvern Instruments) operating at a laser wavelength of 633 nm equipped with a Peltier temperature controller set to 25 °C. Data were collected on 50 μM protein samples in NMR buffer in 3 x 3 mm cuvettes. Using the Malvern DTS software, mean hydrodynamic diameters were calculated from three replicates in the volume-weighted mode.

### Dityrosine fluorescence detection

Steady-state fluorescence detection of dityrosine crosslinks was performed on a Jasco J-720 fluorimeter at 5 μM sample concentrations, in 3 mm cuvettes at 25 °C. Excitation was set to 315 nm and emission spectra were recorded from 325 to 550 nm (λ). The dityrosine-specific emission signal was monitored at 410 nm according to (66).

### Western blotting

SDS-PAGE separated proteins were transferred onto nitrocellulose membranes and blocked in 5% non-fat milk dissolved in PBS supplemented with 0.01% Tween for 1 hour. Membranes were incubated with the following monoclonal primary antibodies at 4° C, overnight. Anti N-terminal αSyn (Abcam, ab51252, 1:1000 dilution), C-terminal αSyn (Santa Cruz, sc-12767, 1:200 dilution), C-terminal αSyn (Santa Cruz, sc-69977, 1:100 dilution), full-length cyt *c* (Abcam, ab13575, 1:500 dilution) and dityrosine (antibodies-online, ABIN361276, 1:100 dilution). Membranes were washed and incubated with anti-mouse or anti-rabbit secondary antibodies (1:5000 dilution) conjugated to horseradish peroxidase (HRP). Proteins were visualized by enhanced chemiluminescence (ECL) detection on a ChemiDoc XRS+ imaging system (BioRad).

### Nuclear magnetic resonance (NMR) spectroscopy

NMR experiments were performed on Bruker Avance 600 or 750 MHz spectrometers, equipped with cryogenically-cooled triple resonance ^1^H{^13^C/^15^N} TCI probes and z axis self-shielded gradient coils. 2D ^1^H-^15^N correlation spectra of monomeric uniform ^15^N nitrogen labeled αSyn and HMWAs were acquired using the SOFAST-HMQC pulse sequence on 50 μM samples dissolved in NMR buffer at 10 °C, with 1K and 256 complex points for sweep-widths (SW) of 16 ppm and 28 ppm in ^1^H and ^15^N dimensions, respectively. Spectra were recorded with 32 scans and interscan delays of 60 ms. Processing was done by zero-filling to 4 times the number of complex points, followed by a sine modulated window function multiplication, Fourier transformation (FT) and baseline correction in both dimensions. Diffusion ordered spectroscopy (DOSY) was performed on 50 μM monomeric αSyn and homogeneous HMWA samples in NMR buffer. Diffusion coefficients were obtained by fitting intensity curves from bipolar pulse-pair longitudinal-eddy-current delay (BPPLED) experiments with an array of linearly shifted gradient power. A series of 28 1D NMR spectra were collected as a function of gradient amplitude. Intensities were measured by integrating signals from 1D proton (^1^H) NMR spectra between 0.75 ppm and 3.2 ppm. Monomeric αSyn and its hydrodynamic radius/diameter (Rh) obtained by DLS was used as the internal reference to determine Rh for HMWAs. ^15^N amide-backbone relaxation data were acquired using standard pulse sequences provided in the Bruker Topspin library. Spectra used for longitudinal relaxation T_1_ (1/R_1_) analysis were collected using the following delay times (in ms): 12, 52, 102, 152, 202, 302, 402, 602, 902, 2002, 5002. T_2_ data (1/R_2_) were measured using a pulse sequence employing a CPMG pulse train with the following delays (in ms): 6, 10, 18, 26, 34, 42, 82, 122, 162, 202, 242, 322 and 462. Duplicate spectra were collected for several time points to estimate uncertainty. To determine R_1_ and R_2_ relaxation rates resonance signal intensities were extracted and fit as a function of the relaxation delay time using CCPNmr Analysis 2.1.5. To obtain steady-state hetero-nuclear (het) NOE values, we calculated ratios of peak intensities in paired spectra collected with and without an initial 4 s period of proton saturation during the 5 s recycling delay. For relaxation experiments, the concentrations of αSyn and HMWA aggregates were 50 μM. Resolution settings and processing were the same as for 2D SOFAST-HMQC experiments. 8, 16 and 32 transients were used for T1, T2 and hetNOE experiments, respectively. To determine residue-resolved NMR signal intensity ratios, only cross-peaks of well-resolved αSyn residues were used. Absolute HMWA signal intensities (I) were divided by corresponding signals of monomeric αSyn (I_0_) and I/I_0_ ratios were plotted against the protein sequence. Acquisition, processing, and spectral analyses were carried out in TOPSPIN 2.1, iNMR 3.6.3 and CCPNmr 2.1.5.

### Photo-induced cross-linking of unmodified proteins (PICUP)

PICUP HMWAs were generated by reacting 300 μM of ^15^N isotope-labeled αSyn with 2 mM ammonium persulfate (APS) and 150 μM of tris(2,2′-bipyridyl)-dichloro-ruthenium(II)-hexahydrate, Ru(BPY)_3_ to photo-oxidize Ru^2+^ to Ru^3+^ by irradiation via a fiber optic cold-light source (Schott KL1500) for 1 minute. 300 mM DTT was added to stop the reaction. HMWAs were purified by SEC on a Superdex-75 column to separate aggregates from APS, Ru(BPY)_3_ and DTT, pooled and concentrated. 50 μM aliquots of PICUP HMWAs were analyzed by SDS-PAGE and NMR spectroscopy.

### Mass spectrometry (MS)

PICUP HMWAs (20 µg) were digested (protein concentration of mg/mL in a 20 mM HEPES, 20 mM NaCl, 5 mM MgCl_2_ buffer) using the endoprotease Glu-C. HMWAs were first reduced by adding 5 mM dithiothreitol (DTT), with incubation for 30 min at 60 °C, followed by alkylation with 15 mM iodoacetamide, with incubation in the dark for 15 min at 20 °C. Glu-C was then added at a protease-to-protein ratio of 1:13 (w/w) and digestion allowed to proceed for 18 h at 37 °C. The resulting digest was acidified to pH 2 with trifluoroacetic acid and desalted using C18 StageTips before analysis on the mass spectrometer. Peptides were loaded (at a flow rate of 0.5 µl min^-1^) directly on a spray analytical column (75 µm inner diameter, 8 µm opening, 250 mm length; New Ojectives, Woburn, MA) packed with C18 material (ReproSil-Pur C18-AQ 3 µm) using an air pressure pump (Proxeon Biosystems). Two gradients (at a constant flow rate of 0.3 µl min^-1^) were used for chromatographic separation (A: 0.1% formic acid in water, B: 0.1% formic acid in 80/20 acetonitrile:water): Gradient 1 (increase over 95 mins) 1% B (0-24 min), 1-5% B (24-25 min), 5-32% B (25-107 min), 32-35% B (107-114 min), 35-85% B (114-119 min); Gradient 2 (increase over 135 mins) 1% B (0-24 min), 1-5% B (24-25 min), 5-32% B (25-144 min), 32-35% B (144-154 min), 35-85% B (154-159 min). Peptides were sprayed directly into a hybrid linear ion trap – Orbitrap mass spectrometer (LTQ-Orbitrap Velos) and analyzed using a “high/high” acquisition strategy, detecting peptides and at high resolution in the Orbitrap and analyzing fragmentation products also in the Orbitrap. Precursor scan (MS) spectra were acquired in the data-dependent mode, detecting in the Orbitrap at 100 000 resolution. The eight most intense triply charged or higher precursors for each acquisition cycle, were isolated with a 2 Th *m/z* window and fragmented in the ion trap with collision-induced dissociation (CID) at a normalized collision energy of 35. Subsequent product (MS2) fragmentation spectra were then recorded in the Orbitrap at 7500 resolution. A dynamic exclusion window (with single repeat count) of 60 s was applied and automatic gain control targets were set to 1⨯ 10^6^ (precursor scan) and 1 x 10^5^ (product scan). Raw files for crosslinking searches were processed using MaxQuant software (v. 1.2.2.5) using default parameters, except for the setting “Top MS/MS peaks per 100 Da”, which was set to 100. Peak lists were searched against the sequence of αSyn using Xi software (v. 1.7.5.1). The following search parameters were applied: MS accuracy, 6 ppm; MS/MS accuracy, 20 ppm; missing mono-isotopic peaks, 2; variable modifications, carbamidomethylation (Cys) and oxidation (Met); enzyme, V8, with a maximum of four missed cleavages allowed; crosslinker modification mass (Tyr-Tyr), −2.015650 Da. Matched crosslinks were mapped to the primary protein structure using xiNET.

### RCSN3 cell lysates, DEPC treatment and H_2_O_2_ quantification

RCSN3 cells (rat cortical *substantia nigra* neurons) were grown in DMEM HAM F12 medium (PAA) supplemented with 12.5% fetal calf serum (FCS) and 1% penicillin/streptomycin to confluency in 175 cm^2^ flasks, in a 5% CO_2_ humidified environment at 37 °C. Following medium removal and washing with PBS, 100 μL of chemical lysis buffer (50 mM Tris-HCl pH 7.4, 0.1% Triton X, 0.25% Na-deoxycholate, 150 mM NaCl, 1 mM NaF, no reducing agents) was added to each flask. Cells were detached with a plastic scraper and collected lysates were centrifuged at 16.000 x g for 30 min. Total protein concentrations of soluble fractions were determined with a Bradford assay (BioRad) and adjusted to 3 mg/mL with NMR buffer. For inactivation with diethyl-pyrocarbonate (DEPC, Sigma) concentrated stock solutions were prepared in 100% ethanol. DEPC was added to RCSN3 lysates to final concentrations of 50 mM and incubated for 22 h at 25 °C. αSyn (50 μM) and cyt *c* (50 μM) were mixed with native and DEPC-inactivated RCSN3 lysate aliquots and 0.5, 10, 50 and 500 mM of H_2_O_2_ (input concentrations). To determine amounts of remaining H_2_O_2_ in native and DEPC-inactivated cell lysates, a colorimetric peroxide detection kit was employed (AssayDesigns, Stressgen).

### Small unilamellar vesicles (SUVs)

Small unilamellar vesicles (SUVs) were prepared from commercial pig-brain polar lipid extract (Avanti) as reported (38). The lipid powder was dissolved in NMR buffer at a concentration of 16 mg/mL (aprox. 20 mM assuming an average lipid mass of 800 Da) and vortexed at room temperature for 30 min. The solution was frozen and thawed 5 times on dry-ice and a 37 °C water-bath, sonicated for 20 min at 4 °C using 30% sonicator output power (Brandelin Sonoplus) and centrifuged at 16,800 × g for 10 min to remove remnant insoluble material. The resulting average vesicle size was 60 nm as determined by DLS. For the binding experiments we added 50 μM of ^15^N-isotopically enriched αSyn or HMWA to SUV solutions (final concentration ∼ 16 mM after αSyn or HMWA addition). NMR spectra were recorded at 30 °C.

### Thioflavin T (ThT) aggregation assay

Kinetic aggregation assays were performed with 100 μL aliquots of 50 μM monomeric αSyn (in 100 mM Na-phosphate, 10 mM NaCl, pH 7.2), or monomeric αSyn in the presence of 5 μM HMWAs (10%), and 20 µM Thioflavin T (ThT) in low-binding 96-well plates (Corning), shaken at 200 rpm with one 2 mm glass bead per well, at 37 °C. Samples were shaken continuously for 2 h before ThT fluorescence emission was recorded at 485 nm (excitation at 440 nm) on a plate reader (InfinitE M200). Samples were set up in triplicates. Data points in **Figure 4D** correspond to the mean value, error bars denote standard errors of the mean (SEM).

### MTT metabolic assay

PC12 (rat pheochromocytoma, American Type Culture Collection) cells were cultured in DMEM medium (Gibco BRL) supplemented with 5% FBS, 10% horse serum and 3 mM glutamine. Cells were plated at a density of 10^3^ cells per well in 96-well plates coated with polylysine in 90 μL of fresh medium in a 5% CO_2_ humidified environment at 37 °C. After 24 h, aliquots of αSyn protofibrils, pure HMWAs or αSyn -HMWA aggregation mixtures were added at concentrations of 0.03, 0.06, 0.13, 0.25, 0.5 and 1 μM, in three replicates. Cells were incubated for 3 days at 37 °C. Cytotoxicity was measured using a 3-(4,5-dimethylthiazol-2-yl)-2,5-diphenyltetrazoliumbromide (MTT) assay kit (Promega) by determining Formazan absorbance at 590 nm on a Tecan Safire automated plate reader. Absorbance values obtained from samples containing aggregated or fibrillar species and from untreated cells were averaged and used to calculate cell viability. Viability was expressed in percentage compared to untreated cells (100%). Error bars indicate SEM values.

## Supplementary Figure Legends

**Figure S1: Mass spectrometry and NMR analysis of PICUP HMWAs. (A)** Representative fragmentation spectra of αSyn dityrosine peptide-pairs following photo-induced crosslinking (PICUP) and proteolytic digestion. Matched αSyn fragments were identified via automated database searches using the Xi software package. **(B)** Overlay of 2D ^1^H-^15^N NMR spectra of ^15^N isotope-labeled, N-terminally acetylated, monomeric αSyn (50 μM, black) and of PICUP-HMWAs (50 μM, red). Resonance signals of N-terminal αSyn residues 1-10 are shown in blue. HMWA resonances broadened beyond detection are labeled, NMR signals of Y39, Y125, Y133 and Y136 are highlighted in grey. NMR spectra were recorded at 283 K (10 °C).

**Figure S2: HMWAs in crowded environments. (A)** Overlay of 2D ^1^H-^15^N NMR spectra of ^15^N isotope-labeled, N-terminally acetylated, monomeric αSyn (50 μM) in the presence of 200 mg/mL BSA (black). N-terminal αSyn residues 1-10 in the reference spectrum are broadened beyond detection due to transient interactions with BSA. Y39 signal intensity of monomeric αSyn is attenuated for the same reason. The corresponding *in situ* NMR spectrum of HMWAs (50 μM) formed in the presence of BSA is overlaid in red. Broadened HMWA resonances are labeled, NMR signals of Y39, Y125, Y133 and Y136 are highlighted in grey. **(B)** Western blot analysis of RCSN3 lysate recovery times following DEPC treatment to determine compound decomposition. After incubation with DEPC, αSyn (50 μM) and H_2_O_2_ (10 mM) were added to lysates after 2 h, 4 h, 6 h and 22 h. Residual DEPC activity was monitored via peroxidative crosslinking of αSyn. **(C)** Overlay of 2D ^1^H-^15^N NMR spectra of ^15^N isotope-labeled, N-terminally acetylated, monomeric αSyn (50 μM) in DEPC-treated RCSN3 lysate (3 mg/mL total protein concentration, black) and *in situ* NMR spectrum of HMWAs (50 μM) formed after addition of cyt *c* (50 μM) and 50 mM H_2_O_2_ (red). HMWA resonances broadened beyond detection are labeled. Chemical shift changes of N-terminal αSyn residues 1-10 are shown in blue. NMR signals Y39, Y125, Y133 and Y136 are highlighted in grey. All NMR spectra were recorded at 283 K (10 °C).

**Figure S3: αSyn and HMWA SUV-binding. (A)** Overlay of 2D ^1^H-^15^N NMR spectra of ^15^N isotope-labeled, N-terminally acetylated, monomeric αSyn (50 μM, black) and in the presence of small unilamellar vesicles (SUV) prepared from pig-brain polar lipid extract. Non-broadened NMR signals of detectable C-terminal residues 106-140 are labeled. Y125, Y133 and Y136 are highlighted in grey. **(B)** Overlay of 2D ^1^H-^15^N NMR spectra of ^15^N isotope-labeled HMWAs (50 μM, black) and upon binding to SUVs (red). Non-broadened NMR signals are labeled. Spectra were recorded at 303 K (30 °C).

