## Supplementary figures and images for "Megadalton-sized dityrosine aggregates of α-synuclein retain high degrees of structural disorder and internal dynamics"

### Supplementary Figures

#### Figure S1

Verzini *et al.*

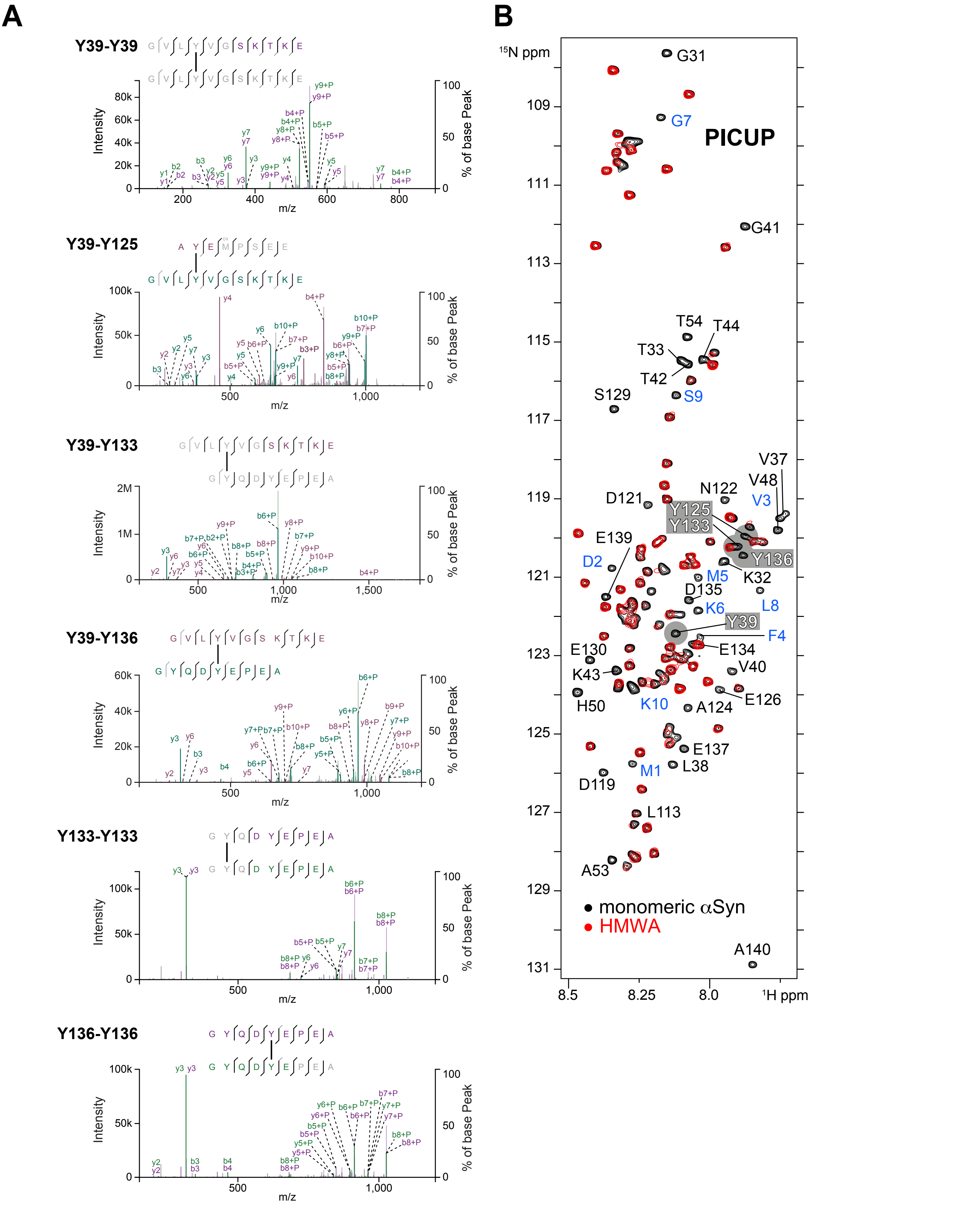

#### Figure S2

Verzini *et al.*

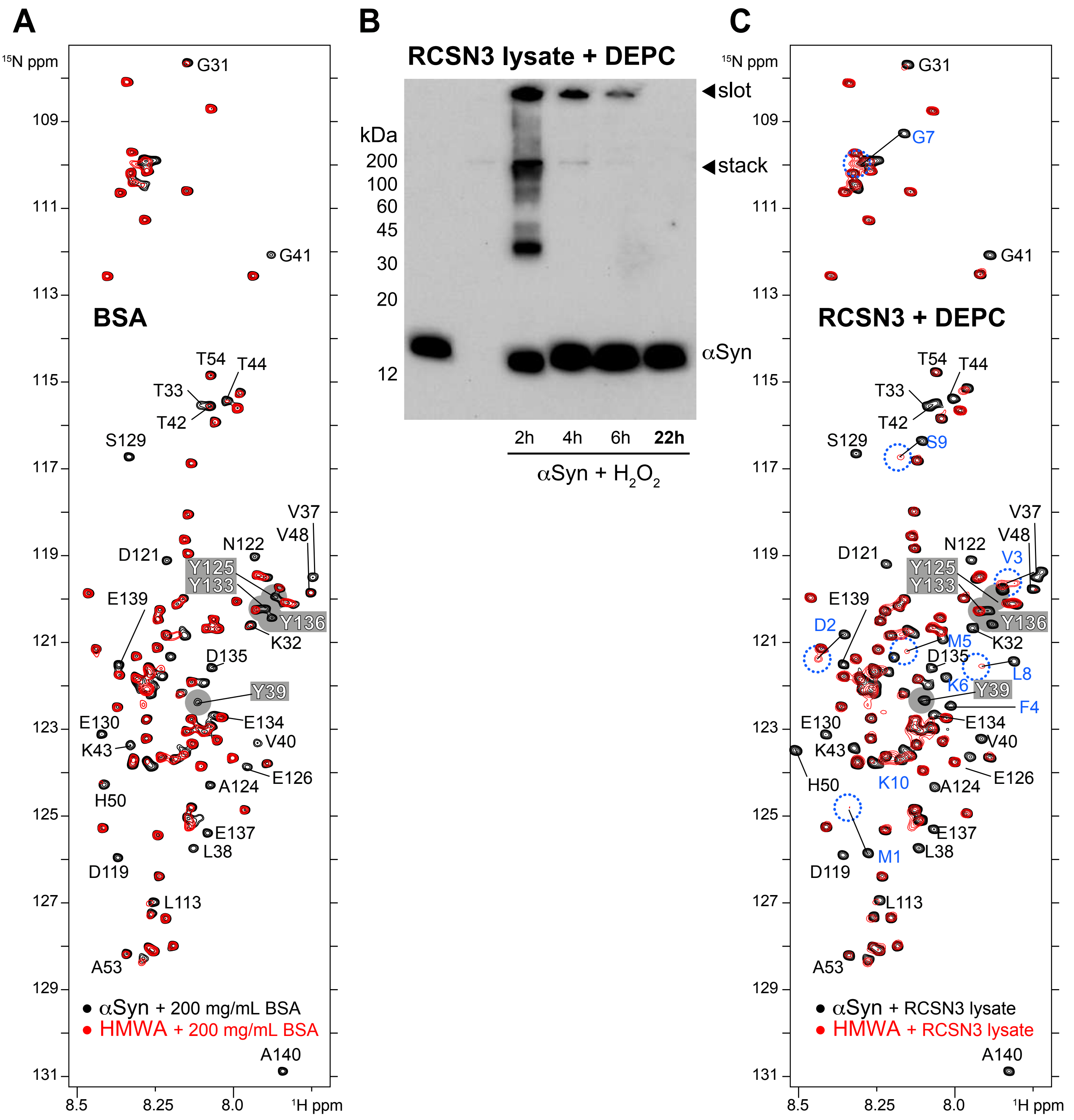

#### Figure S3

Verzini *et al.*

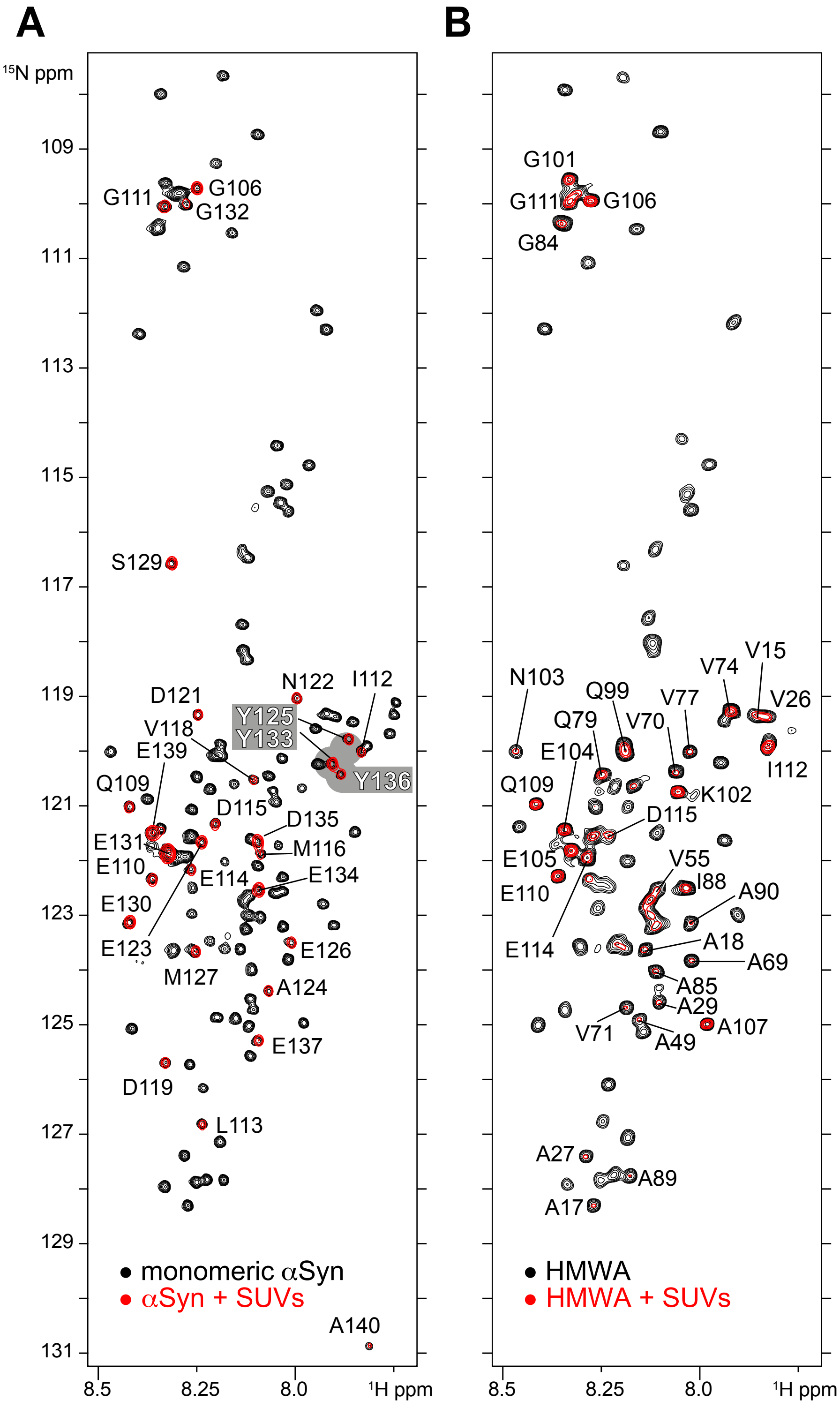
